# Ornamental radiance as a defence against mimicry at the monogamy-polygyny interface

**DOI:** 10.1101/2024.03.17.585419

**Authors:** Andrew Crompton

## Abstract

Ornament evolves though sexual selection when both monogamy or polygyny are equally viable. In this special situation monogamous females will be be able to reject unwanted polygynous suitors only if monogamous males are able distinguish themselves with a sign of fidelity. Since the most sexually advantageous trait value for any male to possess is the one that matches the mean female preference, any signs so created will persist when either monogamy or polygyny subsequently becomes a dominant strategy. Species will therefore accumulate traces of each passage through the monogamous-polygynous boundary which, in birds, can lead, among other outcomes, to a collage of patches of coloured plumage. If a species lives at the interface long enough a polygynous bird might be able to copy the sign of fidelity. One defence against this aggressive mimicry will be to evolve to a sign that is costly or difficult to copy. This process is modelled mathematically showing that if variation of female preference is increased the male phenotype will alter in unpredictable ways that defy a mimic, at a cost of possibly being maladapted. This causes an ornamental radiance that matches ideas of beauty in the human world.

## Introduction

If male ornament is the result of female sexual selection what do females gain by desiring something that is burdensome or dangerous for their male offspring to possess? Standard theory is divided between ornament being a sign of good genes and a non-adaptive explanations, as reviewed in Kirkpatrick and Ryan (1). Since an assessment of male quality can be, and routinely is, made in most species without using ornament a satisfactory theory ought to at least explain why excessive ornament arises in some circumstances and not in others. It seems to lack a purpose sufficient to justify what is invested in it, all of which suggests that there is some factor missing in our understanding of what ornament does. If it is a sign, what distinction does it make?

These differences are resolved in Charles Darwin’s idea that animals both sense and desire beauty, but his account can only be accepted if beauty is shown to be a proxy for something that made offspring more viable, at present cannot be done, Prum, (2). I observe that beauty, as described in aesthetics, is a quality with odd properties, see Hyman, (3) for a summary. It is not something that is recognised after considered judgment, like proving structural stability or reaching verdict in court. Nor can it be measured except by observing how a creature reacts to it, or by asking a person to rate it. and reasons for the rating, if they can be given at all, are usually vague or negative, such as not ugly. In particular, beauty cannot be produced by an an algorithm, in short, it appears as something that cannot be described. Suppose we accept this; then an interesting question arises. In what circumstances might an animal want to be hard to describe? This suggests that beauty might be a form of camouflage.

In what follows some idea of how mating preferences function genetically is needed. I will use Kirkpatrick and Ryan’s null model of intersexual selection. Given genetic variation for a display trait and a mating preference, the most sexually advantageous trait value for any male to possess is the one that matches the mean female preference.

Specifically, if a preference for one type of male is established by some mechanism, then alternative preferences for males that survive either better or worse will not be favoured. Any equilibrium is stable. It follows that it is how the mean female preference moves from one point of equilibrium to another that needs an explanation, not why it is has such and such a value. It value at any particular time is given by its history. Kirkpatrick and Ryan describe three kinds of evolutionary force that can move the female preference: direct selection of preferences; indirect selection in a runaway process, and indirect selection by the parasite mechanism. It is the first kind of force that will be used here. The idea is that female preference shifts because their fecundity is affected by a conflict between monogamous and polygynous mates. This is resolved by the evolution of a mark that distinguishes monogamously inclined males from their polygynous kin. The female accepts that the male with mark might carry a handicap because her alternative is not to reproduce at all. A mathematical model that supports this idea follows. In principle it ought to apply to all cases of sexually selected ornament and behaviour, but for simplicity of expression it is presented here in terms of birds possessing features such as feathers with such and such a length and intensity of colour.

Most birds are monogamous since it usually takes a pair to tend a nest, but polygyny can also be viable, the point is that one or other strategy will be dominant, so a bird following the wrong path will be eliminated by natural selection. Which strategy is dominant depends on circumstances, but when those circumstances change there will be a moment, perhaps a long moment, when both are equally viable. In this intermediate zone, where natural selection is not decisive, the payoff matrix for different couplings of monogamous (M) and polygynous (P) parents is as shown here. The values give the probability of offspring, other factors being equally favourable.

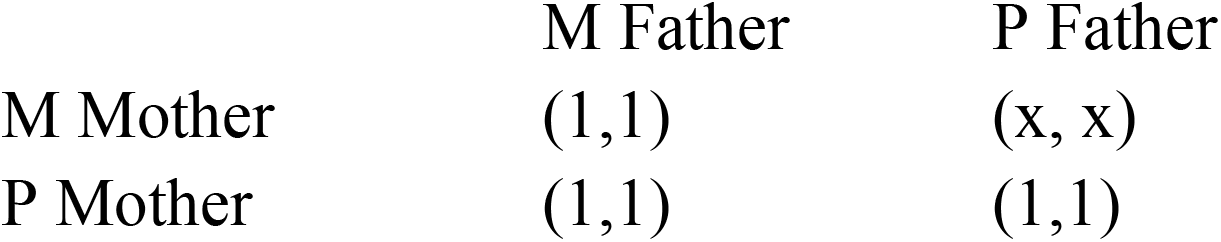

|  | M Father | P Father |
| --- | --- | --- |
| M Mother | (1,1) | (x, x) |
| P Mother | (1,1) | (1,1) |

The value of x has a different significance to the mother and father because of the difference in their gamete size. If x is less than one but not actually zero it profits a polygynous male to mate with a monogamous female since he can find other opportunities to reproduce, but the monogamously inclined female will be at a disadvantage. One possible outcome is that she will be eliminated by natural selection the instant polygyny becomes viable. Another is that her offspring will survive if she can find a way to distinguish monogamous and polygynous suitors. Even if no visible difference exists between them she might be able to elicit a difference. Suppose that under the stress of finding a suitable mate, she broadens the mean female preferences slightly, from average to something close to average. In that case a monogamously inclined male might through natural variation and pleiotropy, evolve a sign, such as a song, or a dance, or some visible difference. For the sake of argument let us imagine he grows a red feather. Female offspring seeing father in the nest, would learn the sign of a mate that plays fair, and in their turn look for a red feathered mate. There is no reason for such a sign to be excessive or maladaptive, other than than it makes the males slightly conspicuous. If circumstances change so that either monogamy or polygyny becomes dominant the sign becomes redundant but it will persist. This is because, as Kirkpatrick demonstrates, there is a line of evolutionary equilibria relating the average female mating preference to the average male trait in the population. Having reached a point on this line, there will be no further shift in the female preferences, unless, for instance it affects their fecundity by making their mates vulnerable to predators, and we will assume that the red feather is not so conspicuous that it is dangerous. Thus crossing between monogamy and polygyny will cause speciation, in this example a split into monogamous-red and polygynous-plain. A species that oscillates across the intermediate zone will accumulate signs. This might explain birds, such as finches, which seem to be of more or less one species, but come in variants with many patches of colour. These are relics of isolated populations that have crossed the zone and made their individual sequence of pleiotropic responses that trace their zig-zag passage across the zone.

This speciation is not, however, the only possible outcome. Suppose a population of birds spends many generations in the ambiguous zone, and suppose that x is greater than zero. In these circumstances a strange contest can take place if a polygamous bird is able to mimic the sign of monogamy. Fair birds try to create a distinction, and cheating birds try to cross it, lying camouflaged among the fair birds like Batesian mimics among truly poisonous animals. If the mean male traits of desirable males are being mimicked by a parasite what options are open to a monogamous female? She must induce males to make a sign that cannot be copied. Strictly speaking, this might be impossible, what one bird can do so can another. The best that can be achieved is something hard to copy. In the human world such things include keys and passwords that depend upon secrecy, but birds have no way to share a secret sign.

Money is made hard to copy by being made of scarce precious metals, or as with printed money by using designs too complicated to copy in a finite time, and which are regularly changed. A similar strategy would make the sign of fidelity costly, or to be more precise it would try to ensure that a failed copy of the sign is too costly to be worth trying. It follows that any sign that monogamous birds use to identify one other cannot be fixed and must change faster that the impersonator can respond; but if it changes how can its meaning be established? All these strategies depend upon some agreement among users to fix authenticity and seem unavailable to an animal. This is the puzzle that evolution has to solve.

One way do it might be to base the sign on a property possessed by the whole group, that can be measured by any bird, and which can alter. Relating the sign to an average bird is one way to do this. Thus problem has the simple solution, in fact it is the same strategy used to induce monogamous males to distinguish themselves from their polygynous kin. Females respond to stress of not being able to identify a reliable mate by allowing even more variety in what they consider acceptable. Their mean preference broadens. The effects of this change are unexpected and dramatic.

If the female allows a single trait, such as the red feather, to increase then its average will take a bounded random walk between limits of no signal up to point where it becomes a handicap. The mimic can follow, but at a cost of reducing its attractiveness to other polygynous females who are indifferent to ornament, as well bearing as the handicap that will make mimics less viable. A single feature with natural variation will not be sufficient to differentiate birds safely. Better is a combination of features change in irregular ways faster than an imposter can respond. Let us call a male that matches the mean female preference the most beautiful bird. If this mean involves comparing several traits then the idea of an average becomes harder to judge. If we imagine a sum of averages then there are more ways to be beautiful, since one feature can compensate another. When his process is modelled one finds that what is beautiful shifts unpredictably from generation to generation. This makes the mimic’s work harder as there is a moving target.

Tiny differences between multiple features might be hard to detect and go some way to explaining the peculiar behaviour of female birds-of-paradise. They are connoisseurs of male display, one study found they watched male displays for six hours per day, and nearly always say no, requiring about fifty hours of observation per mating event, leading to only 1.2 matings per season for the most successful males. Most males did not mate at all, (Pruett-Jones). Despite the flamboyance females are looking for something that is hard to see. I suggest that they are establishing a sense of the average bird then choosing one as close as possible if he is not available. If the mean female preference is not innate but measured. What is most beautiful bird changes more or less randomly and becomes hard to mimic.

Beauty does not mean above average features, difference can be in either direction, pressure on the phenotype is to move in unpredictable ways to foil the mimic. Only one bird however is the most beautiful at any one time, but since monogamy removes beautiful fathers from the pool, so the range of acceptable, good enough mates must be also broad and this gives some margin of error for the mimic. This imposes a limit on the number of safe acceptable mates.

Features follow a bounded pseudo-random walk. If they become fixed or go to nothing the game is over. The are not truly random because they are weakly coupled to each other. It is in assessing the most beautiful bird that the features interact. A sudden leap in the value of a feature in one bird will prevent it from being beautiful but will increase the average, and can passed down the female line though its sisters. They can range randomly in the positive direction they will become maladaptive. Excessive features are accidental, they are not making, as it were, making a big statement, but only the result of a random movement in feature space to avoid being mimicked. All the ornament delivers a single bit of almost honest information.

## Materials and methods

The process was modelled by representing birds as a set of numbers corresponding to values given to features. These were imagined as representing the colour intensity or length of feathers although they ought to be equally applicable to other modes of making a sign such as sound or movement.

A creature’s ability increase the value of a feature is limited by natural selection, some features can hardly be altered, but a phenologically plastic creature will have potential to make differences in plumage, or in song at little cost to its viability. There will have lower limit, either nothing, in the case of a difference in colour, or one if a change in ratio of some size. In what follows, for the sake of a simplicity I will take the initial value of features to be one. Small downward movement is allowed, (representing a short tail for instance). Feature were given an upper and lower limit corresponding to a functional minimum and being a handicap.

The model, a vba module, had m male and m female birds b_i_ (i = 1,2, . . m), each with n features expressed in the nubile male bird, latent and inheritable in the female, f_j_ (j = 1,2, . . n). Each year the male birds are inspected and the average value of each feature measured, how this changes year by year in shown in the figures. The values of f_ave_, what an ornithologist sees, were were plotted against time for various values of parameters m, n, p, h, etc.

How far each bird is from average for each feature is measured as (f_j_ - f_ave_)/ f_ave_ in accordance with the Weber-Fechner law in which the perceived difference varied with the size of the stimulus. The sum of these perceived differences is made for each bird, ∑ f_j_ - mean j = 1,2, . . n, i = 1,2, . . m, so the male birds are ranked according to how far they are above or below average. The bird closest to average is called most beautiful. If p couples mate per year then the p males whose rank is centred on the most beautiful mate with p females chosen randomly, (e.g., if m = 61 and nine couples form each year and the most beautiful is ranked 30 then males ranked 26 to 34 become fathers.) Their offspring are a pair of male and female birds that replace the oldest birds. All birds live the same number of years so the population is constant. New birds have feature values chosen from a normal distribution with the same standard deviation as the whole population and a mean of (1-h). population mean + h. mid-parent value, where h is a hereditability factor between 0 and 1. When h = 0 then there is no parental influence on offspring, when h = 1 offspring copy their midparent’s values.

All the figures here show the model with 67 birds with 10 features, an attempt to model the lek of a bird of paradise, *Lawes Parotia* observed by Pruett-Jones, (4), in which 67 different females visited the lek and 9 different males mated each season. Pruett-Jones observed that females were monogamous during each season, noting that the loss of a clutch did not alter females choice of mates. The model starts with features for each bird set at an average value of 1 with a standard deviation of 0.1. The model’s parameters were varied to show how average feature values for a population of birds were affected over time, typically 1000 generations.

## Results

If the features settle on fixed values then a mimic can copy those values and the game ends. The question is, under what conditions do the feature values change unpredictably? The range of parameter values for which this happens is fairly wide. Figures 1 shows results where this is the case.

**Figure 1.**
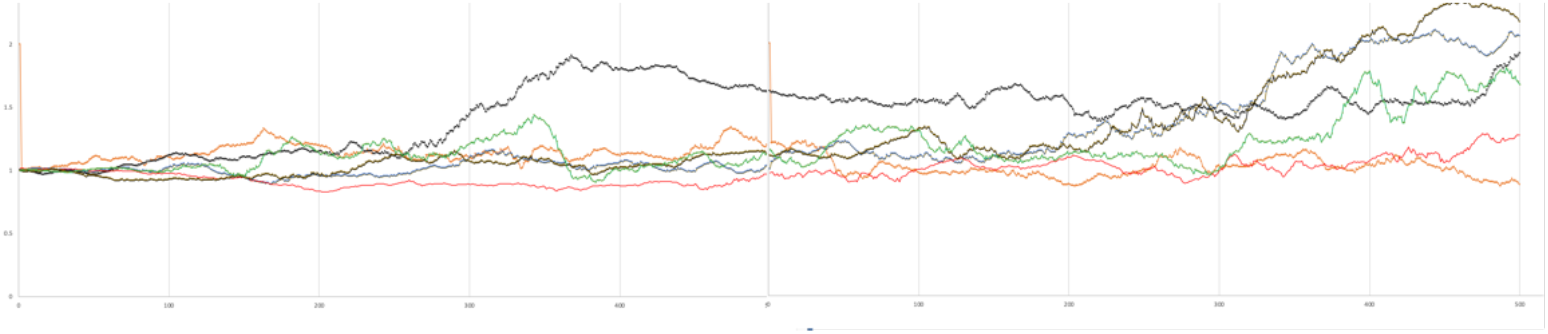
Evolution of average feature values for 67 birds, 10 features, h = 0.5, upper limit of 2.0, lower limit 0.6, with 9 mating pairs per season, over 1000 generations. This chart is a typical outcome of the model.

i. It does not happen with a single feature which will rise and fall gradually presenting a wide target to a mimic.
ii. When the hereditability factor was set at zero then, unsurprisingly, features did not change but preserved their small initial variation. Only when h was increased above about 0.3 did the features began to wander in unpredictable ways.
iii. The effects of females varying their preferences were modelled by changing the upper and lower limits of features. When they were close together, e.g. lower limit of 0.8, upper limit of of 2, then the variation in features was small. As the upper limit was increased then average feature value also increased. When this happened the values did not simply rise but varied in a chaotic way as seen in Figures 1 and 2.

**Figure 2.**
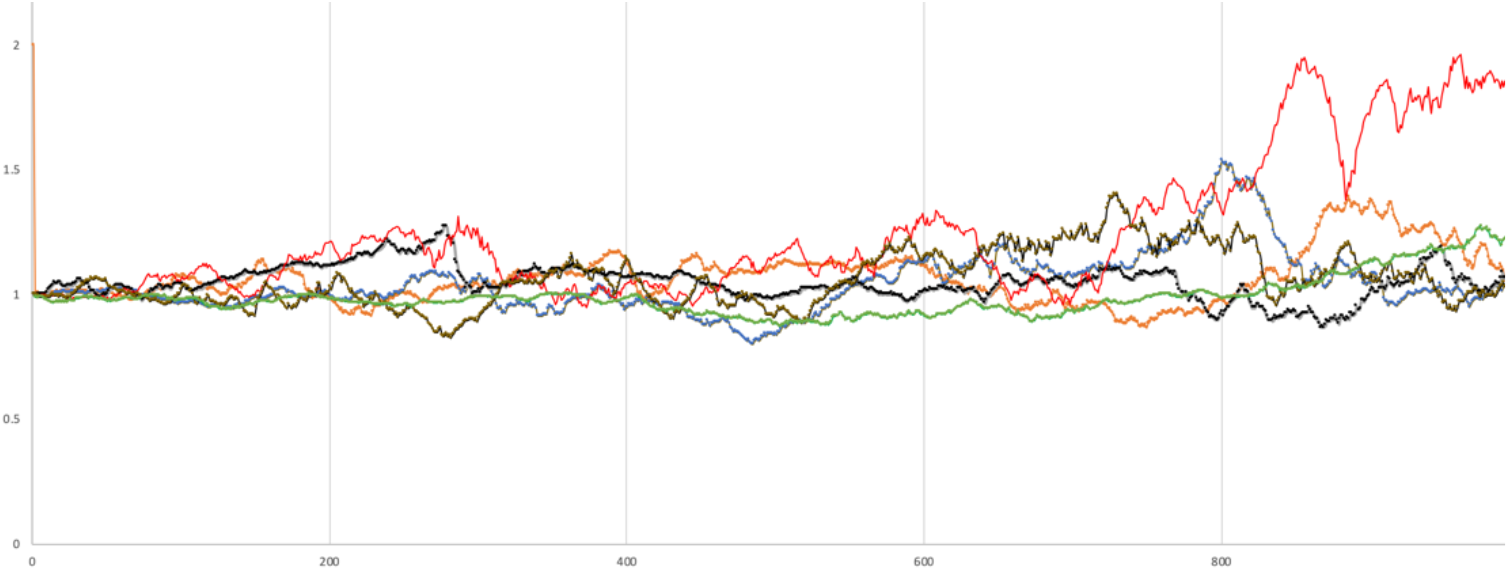
Average feature values for bird over 1000 generations, values as figure 1 but with upper limit of 4.0, causing more variety.
iv. Increasing the number of birds and increasing the number of features made the average value of features change more slowly, as if the system had more inertia, but it did not change the pattern described above.
v. If a mimic, (or a beautiful male that switched to being polygynous) succeeds in mating its genes will spread through natural drift so long as its offspring are also beautiful, it advantage last only so long as beauty is hereditary, but this might not be case. The model set the breeding life of a bird at ten years, in which case not all birds had offspring. If this restriction was removed so that birds did not die, then I observe that all birds eventually had offspring, and at about the same frequency, so the state of being most beautiful moved through the whole population as the same time that the feature strength wandered chaotically. In the model beauty is not be something that can be inherited in a reliable way, rather it is something aleatory. The random drift in features that can be maladaptive. What the model shows is the opposite of convergent evolution, rather is divergent evolution. Crossing the monogamy-polygyny interface causes something general to become more specific like white light becoming coloured by passing through a dispersive medium.
vi. If the ornament is essentially random but endures even after the boundary crossing is complete this explains the uniformity of adult males within each bird of paradise species, and also the divergence between species. Separate groups following the same strategy will reach utterly different solutions. That point that the range of avian plumage needs an explanation, as well its existence was raised by Diamond, 1986 (5).
vii. The circumstances in which this idea of beauty can function are limited. It a group is too large the average will have more inertia allowing mimics a more stable target, moreover females might not be able to measure the average very well. Pruett-Jones found that females spent six hours a day inspecting about twenty males. The point is to absolutely exclude polygamous males then it is necessary that the distinction between nubile and not-nubile be made within the population of monogamous mates, and it here that sexual selection does its work by sacrificing some males, not all of whom can be beautiful at once.

## Discussion

The theory also gives some idea of the conditions necessary for elaborate ornament to arise. To survive the maladaptive possibilities of the radiance the creature should not depend on camouflage to hunt or survive. It needs phenotypic plasticity, and senses capable of recognising features, an exaptation perhaps of those evolved to find fruit or insects. It should find monogamy and polygyny equally viable, or have done so in the recent past, and exist in a society in which a female can assess an average mate, or be able to assemble in leks so this judgement can be made. Moreover the group should be small enough for the average to be able to change faster than a mimic can evolve. These conditions are satisfied in small society of isolated creatures that can fly out of danger, such as a bird in a fractal forest environment.

A speculation: Although these remarks have dealt with visual signs they could equally be applied to signs made by song or dance. Many birds, notably bowerbirds, can copy sounds, including unnatural noises such as saws and camera shutters, as well as any digital recorder. Why might it have been useful to have evolved this ability? Perhaps their ancestors deterred mimics by singing a song that could not be described, and then, due to some neurological breakthrough in the ability to memorise and reproduce sounds the evolutionary arms race was won by the mimic. The ability to copy sounds would of course be retained, the birds, one speculates, then reduced to manufacturing a new uncopiable sign of fidelity out of grasses and found objects as a substitute for a song that since it was no longer indescribable is now lost for ever.

Darwin’s idea that animals can appreciate beauty has some merit if beauty and being average are correlated. There is experimental evidence, for humans at least, that this is indeed the case, (6). Ideas of the beautiful average are at least as old as Durer, who wrote, ‘I hold that the perfection of form of form and beauty is contained in the sum of all men.’ (7) True beauty, according to many thinkers, is better described as close to average, an idea contained in Edmund Burke’s observation that ‘beauty should shun the right line, yet deviate from it insensibly.’(8) Thus we can add indescribable beauty to the list of things that, as a matter of fact, the natural world has reached before humans thought of them.

## References

1 Mark Kirkpatrick & Michael J. Ryan The evolution of mating preferences and the paradox of the lek. Nature 350-7 March 1991 33–38

2 Richard O. Prum, Aesthetic evolution by mate choice: Darwin’s really dangerous idea. Phil. Trans. R. Soc. B (2012) 367, 2253–2265 doi:10.1098/rstb.2011.0285

3 John Hyman Is Beauty In The Eye Of The Beholder? Think Spring 2002 pp. 81–92

4 S. G. & M. A. Pruett-Jones, Sexual Selection Through Female Choice In Lawes’ Parotia A Lek-Mating Bird Of Paradise. Evolution, 44(3), 1990, pp. 486-501]

5 Jared Diamond, Biology of Birds of Paradise and Bowerbirds. Annual Review of Ecology and Systematics,Vol. 17 (1986) 17–37

6 D.I. Perrett, K.A. May, S. Yoshikawa, 1994. Facial shape and judgements od female attractiveness. Nature 368, 239–242 17 March 1994.

7 Jamin Halberstadt and Gillian Rhodes, It’s not just average faces that are attractive: Computer-manipulated averageness makes birds, fish, and automobiles attractive. Psychonomic Bulletin and Review 2003, 10(1), 149–156.

8. Four Books on Human Proportion, Albrecht Durer, trans. by William M. Conway, in Henrietta Midonick, The Treasury of Mathematics Vol 2. Penguin 1965. p.121

8 Edmund Burke, 1759 A Philosophical Enquiry, Oxford: OUP 1990. Section XXVII p.113

